# Spatial transcriptomic profiling of coronary endothelial cells in SARS-CoV-2 myocarditis

**DOI:** 10.1101/2022.09.25.509426

**Authors:** Camilla Margaroli, Paul Benson, Maria G Gastanadui, Chunyan Song, Liliana Viera, Dongqi Xing, J. Michael Wells, Rakesh Patel, Amit Gaggar, Gregory A. Payne

**Affiliations:** Department of Medicine, Division of Pulmonary, Allergy & Critical Care Medicine, University of Alabama at Birmingham, Birmingham, AL, USA; Program in Protease/Matrix Biology, University of Alabama at Birmingham, Birmingham, AL, USA; Department of Pathology, Division of Anatomic Pathology, University of Alabama at Birmingham, Birmingham, AL, USA; Translational Program for Cardiopulmonary Disease, University of Alabama at Birmingham, Birmingham, AL, USA; Lung Health Center, University of Alabama at Birmingham, Birmingham, AL; Medical Service at Birmingham VA Medical Center, Birmingham, AL; Department of Pathology, Division of Molecular & Cellular Pathology, University of Alabama at Birmingham, Birmingham, AL, USA; Center for Free Radical Biology, University of Alabama at Birmingham, Birmingham, AL, USA; Department of Cell, Developmental, and Integrative Biology, University of Alabama at Birmingham, Birmingham, AL, USA; Vascular Biology and Hypertension Program, University of Alabama at Birmingham, Birmingham, AL, USA; Comprehensive Cardiovascular Center, University of Alabama at Birmingham, Birmingham, AL, USA; Department of Medicine, Division of Cardiovascular Disease, University of Alabama at Birmingham, Birmingham, AL, USA

**Keywords:** Spatial transcriptomics, Severe acute respiratory syndrome coronavirus-2 (SARS-CoV-2), Myocarditis, Endothelial

## Abstract

**Objectives:** Our objective was to examine coronary endothelial and myocardial programming in patients with severe COVID-19 utilizing digital spatial transcriptomics.

**Background:** Severe acute respiratory syndrome coronavirus-2 (SARS-CoV-2) has well-established links to thrombotic and cardiovascular events. Endothelial cell infection was initially proposed to initiate vascular events; however, this paradigm has sparked growing controversy. The significance of myocardial infection also remains unclear.

**Methods:** Autopsy-derived cardiac tissue from control (n = 4) and COVID-19 (n = 8) patients underwent spatial transcriptomic profiling to assess differential expression patterns in myocardial and coronary vascular tissue. Our approach enabled transcriptional profiling *in situ* with preserved anatomy and unaltered local SARS-CoV-2 expression. In so doing, we examined the paracrine effect of SARS-CoV-2 infection in cardiac tissue.

**Results:** We observed heterogeneous myocardial infection that tended to colocalize with CD31 positive cells within coronary capillaries. Despite these differences, COVID-19 patients displayed a uniform and unique myocardial transcriptional profile independent of local viral burden. Segmentation of tissues directly infected with SARS-CoV-2 showed unique, pro-inflammatory expression profiles including upregulated mediators of viral antigen presentation and immune regulation. Infected cell types appeared to primarily be capillary endothelial cells as differentially expressed genes included endothelial cell markers. However, there was limited differential expression within the endothelium of larger coronary vessels.

**Conclusions:** Our results highlight altered myocardial programming during severe COVID-19 that may in part be associated with capillary endothelial cells. However, similar patterns were not observed in larger vessels, diminishing endotheliitis and endothelial activation as key drivers of cardiovascular events during COVID-19.

**Condensed Abstract:** SARS-CoV-2 is linked to thrombotic and cardiovascular events; however, the mechanism remains uncertain. Our objective was to examine coronary endothelial and myocardial programming in patients with severe COVID-19 utilizing digital spatial transcriptomics. Autopsy-derived coronary arterial and cardiac tissues from control and COVID-19 patients underwent spatial transcriptomic profiling. Our approach enabled transcriptional profiling *in situ* with preserved anatomy and unaltered local SARS-CoV-2 expression. We observed unique, pro-inflammatory expression profiles among all COVID-19 patients. While heterogeneous viral expression was noted within the tissue, SARS-CoV-2 tended to colocalize with CD31 positive cells within coronary capillaries and was associated with unique expression profiles. Similar patterns were not observed in larger coronary vessels. Our results highlight altered myocardial programming during severe COVID-19 that may in part be associated with capillary endothelial cells. Such results diminish coronary arterial endotheliitis and endothelial activation as key drivers of cardiovascular events during COVID-19 infection.

**LIST OF HIGHLIGHTS:**

1. SARS-CoV-2 has variable expression patterns within the myocardium of COVID-19 patients
2. SARS-CoV-2 infection induces a unique myocardial transcriptional programming independent of local viral burden
3. SARS-CoV-2 myocarditis is predominantly associated with capillaritis, and tissues directly infected with SARS-CoV-2 have unique, pro-inflammatory expression profiles
4. Diffuse endothelial activation of larger coronary vessels was absent, diminishing large artery endotheliitis as a significant contributor to cardiovascular events during COVID-19 infection.

## INTRODUCTION

The unprecedented COVID-19 pandemic caused by severe acute respiratory syndrome coronavirus-2 (SARS-CoV-2) has resulted in immeasurable personal and global loss. With over 50 million confirmed infections, and a death toll eclipsing one million people, the COVID-19 pandemic has caused the deadliest period in U.S. history (data from the Centers for Disease Control and Prevention). Infection with SARS-CoV-2 is characterized by pneumonia, acute lung injury, and subsequent multiorgan failure^1^. Unique from other viral pneumonias, the virulence of SARS-CoV-2 is due, in part, to its ability to induce thrombotic and cardiovascular events^2^. Such complications were speculated to result from direct myocardial injury via viral myocarditis and/or damage to the vascular endothelium^3^. Despite such assertions, we have poor understanding of the molecular mechanisms by which SARS-CoV-2 causes vascular inflammation, cardiovascular collapse, and ultimate death. More importantly, we have little mechanistic evidence explaining why patients with underlying cardiovascular diseases or risk factors appear to suffer the most severe complications.

Disseminated intravascular coagulation^4^ and cardiovascular injury associated with SARS-CoV-2 infection was initially postulated to be a consequence of uncontrolled inflammation and direct endothelial damage induced by SARS-CoV-2 infection^5^. Endothelial expression of the Angiotensin Converting Enzyme 2 (ACE2) receptor for SARS-CoV-2 entry^6, 7^ was thought to mediate endothelial injury. This assertion was supported by initial case series which proposed endothelial cell infection and associated endotheliitis within a wide variety of endothelial cells in COVID-19 patients^3^. Despite such clinical observations, validating this early hypothesis has proven challenging. While animal models utilizing transfected humanized ACE2 have observed SARS-CoV-2 infection of endothelial cells^8^, subsequent translational efforts have failed to observe similar findings. Notably, McCracken *et al*. reported a lack of ACE2 expression and replicative infection by SARS-CoV-2 in human endothelial cells^9^, while more recent autopsy analysis of COVID-19 patients failed to observe coronary endothelial activation^10^. These opposing observations beg the question as to whether direct endothelial mechanisms exist linking COVID-19 to cardiovascular events.

Debate also continues as to whether SARS-CoV-2 causes a classic viral-mediated myocarditis. Acute systolic heart failure of varying severity has been associated with COVID-19 infection, including as many as 10% of hospitalized patients having severe left ventricular dysfunction^11^. However, evidence of direct SARS-CoV-2 cardiomyocyte infection has been variable. Investigators from the COVID-19 tissue atlas^12^ reported little evidence of viral replication in cardiac tissue; however, COVID-19 has been linked with transcriptional alterations in both patient samples^13^ and models of COVID-19 myocarditis^14^. These incongruent findings, in combination with our limited understanding of SARS-CoV-2-induced vascular inflammation, hinders our ability to treat cardiovascular conditions associated with COVID-19. Sadly, despite ongoing COVID-19 vaccine distribution, novel variants are likely to become endemic, posing a long-term danger to at risk populations^15^. Hence, the urgency to discover new diagnostic and therapeutic options against this deadly disease cannot be understated.

Our group recently uncovered differential transcriptional signatures associated with severe SARS-CoV-2 lung injury when compared to classical H1N1 influenza pneumonia^16^. Using a novel spatial transcriptomic platform, we identified robust transcriptional signatures of inflammation, extracellular remodeling, and alternative macrophage activation which had previously not been identified in other SARS-CoV-2 transcriptome studies. These differences extended to the pulmonary vasculature where coagulation pathways were markedly upregulated. These findings are significant, as they implicate potential endothelial-specific differences that may directly contribute to increased cardiovascular complications associated with SARS-CoV-2 infection.

Herein, we report the first spatial transcriptomic analysis of the coronary endothelium from COVID-19 patients. We conducted spatial mapping of the myocardium and coronary vasculature of autopsy specimens collected from patients who died because of severe COVID-19 infection. Our approach enabled us to spatially resolve differential expression patterns of coronary endothelial cells and cardiac tissue without disruption of tissue architecture within preserved patient samples. In so doing, we examined the paracrine effect of SARS-CoV-2 expression on myocardial and coronary endothelial expression profiles.

## METHODS

### Materials and data availability

Anonymized datasets generated during this study will be available on Mendeley Data. Other requests for resources and reagents are available from the corresponding author upon reasonable request. Requests should be directed to and will be fulfilled by Dr. Gregory Payne. This study did not generate new or unique reagents. See the **Major Resources Table** in the Supplemental Materials for detailed experimental reagents.

### Human subjects

The study was approved by an institutional review board (UAB-IRB 300005258, VA-IRB 1573682). Summary demographic and clinical data are presented in **Supplemental Tables S1 and S2**. Autopsy authorization from next of kin included consent for use of the tissue for research. SARS-CoV-2 samples were collected from patients that died due to pneumonia and acute respiratory failure between March and June of 2020. All SARS-CoV-2 participants had chest imaging consistent with findings of pneumonia and positive nucleic acid amplification test (PCR) for SARS-CoV-2 indicating COVID-19 pneumonia. Myocardial specimens were stained for SARS-CoV-2 nucleocapsid and stratified into quartiles based upon relative mean fluorescent intensities (MFI). The bottom quartile (<25% MFI) was defined as SARS-CoV-2 “Low”, while the top quartile (>75% MFI) was defined as SARS-CoV-2 “high”. Separately, four deceased patients without infection or known cardiovascular disease who died before the existence of SARS-CoV-2 were included as control subjects (all died prior to 2020). All participants were grouped according to COVID-19 status and myocardial SARS-CoV-2 expression as discussed.

### Histology and Immunofluorescence

The presence of myocardial infection with SARS-CoV-2 was determined prior to GeoMX digital spatial profiling by immunofluorescence to the viral nucleocapsid as previously described (see **Supplemental Methods** for detailed protocol)^16^. Briefly, anti-SARS-CoV-2 nucleocapsid (GeneTex, GTX135361; 1:500) and anti-CD31 (Abcam, ab9498, 1:200) antibodies were directly conjugated to fluorescent markers (see ***GeoMX digital spatial profiling*** below). Nuclei were counter stained with DAPI (Biolegend, stock 100ng/ml and 1:1000 in PBS working solution). Images were acquired using the Nikon A1R confocal microscope.

### GeoMX digital spatial profiling. Supplemental Figure S1

illustrates the work-flow utilized for digital spatial profiling. Paraffin embedded tissues were processed and analyzed using a combination of fluorescently labeled anti-SARS-CoV-2 nucleocapsid and anti-CD31 antibodies. Anti-SARS-CoV-2 was conjugated to PE / R-Phycoerythrin (Expedeon Lightning-Link R-PE Conjugation Kit / Abcam, ab102918), while anti-CD31 was linked to Alexa Fluor 488 (Abcam, Lighting-Link ab236553). Nuclei were counterstained with SYTO61 (ThermoFisher, S11343, 1:1000 in PBS working solution). Fluorescent antibodies were combined with the GeoMX Cancer Transcriptome and COVID-19 Immune Response Atlas gene sets with custom probes specific for SARS-CoV-2 lung infection and tissue responses (see **Supplemental Table S3** for SARS-CoV-2 related gene list), totaling 1860 genes. Regions of interest (ROIs) were selected based on 1) viral staining, 2) cellular immunofluorescent profile and anatomic features consistent with coronary vascular tissue, and 3) histologic features consistent with myocardial tissue observed in the hematoxylin and eosin (H&E) stained sections. Supplemental **Figure S2** illustrates the distribution of all ROIs for COVID-19 patients. At least 50 cells per ROI were utilized for analyses.

### Quantification

Libraries of the oligo tags collected with the Nanostring GeoMx platform were quantified as previously described^16^. Please see **Supplemental Methods** for detailed experimental protocol.

### Statistical Analysis

Clinical data is expressed as mean ± standard deviation or n (%) unless otherwise indicated. 1-way ANOVA or Chi square test used to measure differences between groups for continuous and categorical values, respectively. For digital spatial profiling, comparison of all groups (e.g., Control, SARS-CoV-2 Low, and SARS-CoV-2 High) was performed by comparing the technical replicates from each biological group. This approach was taken given the heterogeneous vascularity and presence of endothelial ROIs available per myocardial tissue section. Each figure illustrates replicate ROIs for a given patient sample size. Similar statistical analyses were performed per patient (n = 4 patients per group) without any difference to the final conclusions (see results below). Counts were normalized to log2 and statistical comparisons were performed using a mixed linear model with Benjamini-Hochberg correction to account for false discovery rate. P value threshold for differential gene expression was set at p = 0.02 and log2 fold change of 0.4. All details for the statistical analyses and number of replicates can be found in the figure legends. All analyses for the volcano plots can be found in the supplemental Data Set file.

### Independent data access and analysis

Drs. Camilla Margaroli and Gregory Payne had full access to all the data in the study and take responsibility for its integrity and analysis.

## RESULTS

### Cohort Characteristics

Clinical characteristics are presented by group in **Supplemental Table S1**. Briefly, participants were 67±13 years old, 75% male, and 67% reported a prior cardiovascular diagnosis. Participants were severely ill with a PaO2/FiO2 ratio at the time of ICU admission of 210±194 and APACHE II score 19±10. Half of the subjects received systemic corticosteroids during their hospital course and one (8%) received Remdesivir. Acute events that occurred between the time of ICU admission and death are reported in **Supplemental Table S2**. These did not differ by group.

### SARS-CoV-2 infects the myocardium with severe COVID-19

Histologic review by an independent clinical anatomic pathologist of H&E stained myocardium failed to observe inflammatory cell infiltration in either group (**Figure 1A and Supplemental Figure S2**). Given the uncertainty surrounding SARS-CoV-2 myocarditis, we first tested for viral infection via immunofluorescent staining of SARS-CoV-2 nucleocapsid on myocardial tissue sections. Heterogeneous expression of SARS-CoV-2 nucleocapsid was noted within myocardial tissue of COVID-19 patients. When present, however, SARS-CoV-2 nucleocapsid was observed to colocalize with CD31 stained coronary endothelial cells (**Figure 1B**). This observation was most noticeable within the capillary bed adjacent to the myocardium. Based upon these findings, we sub-grouped the COVID-19 cohort into SARS-CoV-2 “Low” or SARS-CoV-2 “high”.

**Figure 1.**
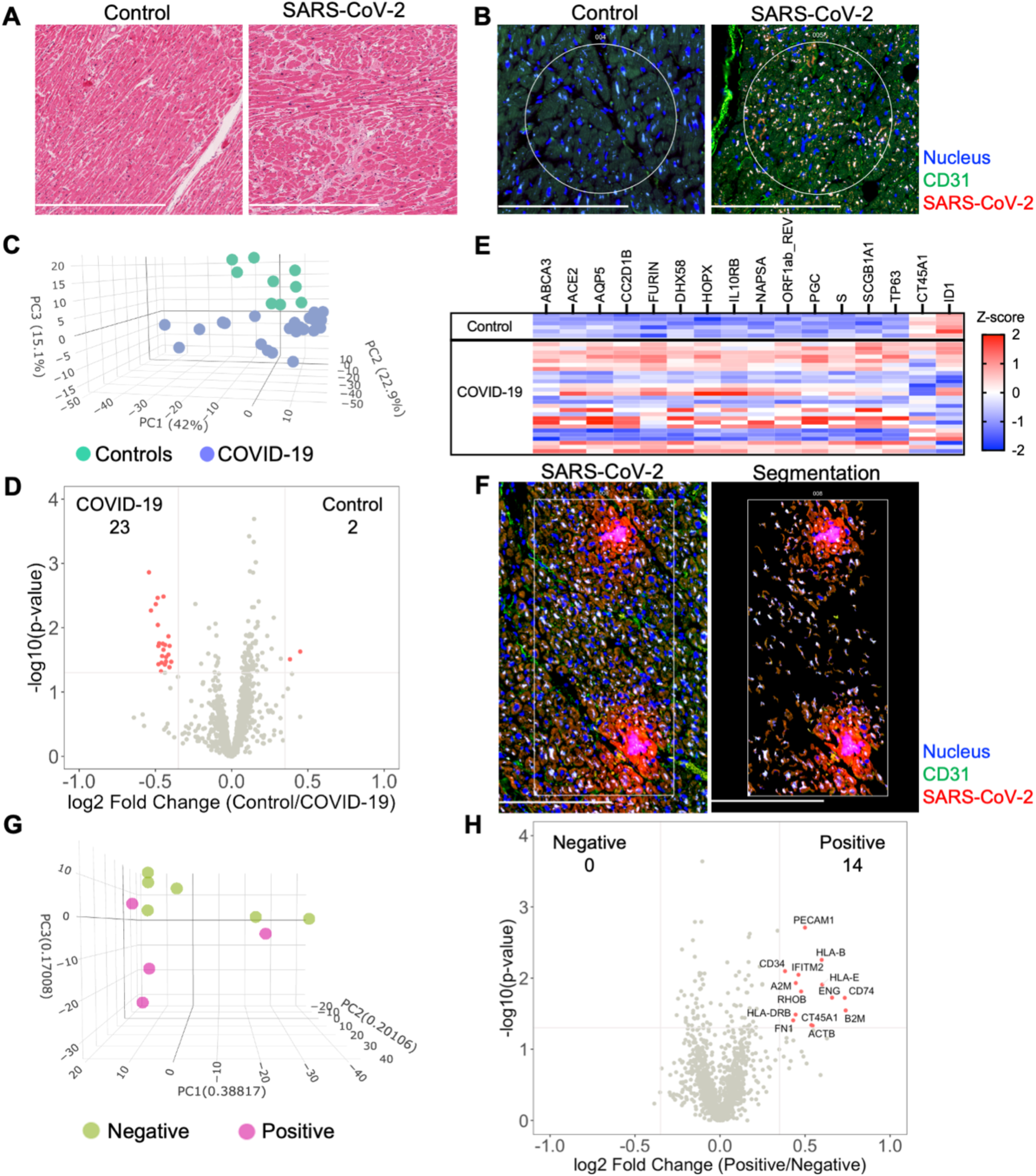
SARS-CoV-2 infects the myocardium and promotes differential expression patterns in patients with severe COVID-19. Histological analysis of the myocardium stained by H&E (A) or by immunofluorescence for SARS-CoV-2 nucleocapsid and CD31 expression (B). Circles highlight representative ROIs. Heterogeneous expression of SARS-CoV-2 was noted among COVID-19 samples. 1454 genes were above the limit of quantification. Compared to controls, PCA of unsupervised data revealed pattern clustering and differential expression within COVID-19 patients (C, data presented per ROI). Gene-expression analysis (D and E) observed 25 differentially expressed genes from myocardial tissues (differential expression was defined as p = 0.02 and log2 fold change of 0.4). Segmented, marker-defined ROIs were selected by SARS-CoV-2 nucleocapsid expression for gene-expression analysis (F, representative image for “positive” ROI). Among COVID-19 patients, PCA of SARS-CoV-2 positive tissue revealed pattern clustering (G), enrichment of endothelial cell markers, and upregulation of key inflammatory genes (H). COVID-19 (n = 8 patients, 25 ROIs) and Control (n = 4 patients, 9 ROIs). Scale bars are 250µm for each image.

### COVID-19 induces a unique global myocardial transcriptomic profile

To investigate similarities and regional differences in COVID-19 myocarditis, we utilized the GeoMX digital spatial profiling platform to sequence specific ROIs from myocardial tissues (see **Supplemental Figure S1** for experimental workflow). Our approach enabled the selection of unique ROIs based upon anatomy and SARS-CoV-2 tissue expression (**Supplemental Figures S1 and S2**) from control and COVID-19 patient samples. Utilizing an RNA *in situ* hybridization gene platform, we initially examined global transcriptional changes in myocardial tissue associated with COVID-19. Among the 1860 genes tested, 1454 genes were above the limit of quantification. Compared to controls, principal component analysis (PCA) of the unsupervised data revealed significant pattern clustering and distinct variation in transcriptional programming among COVID-19 patient samples (**Figure 1C**). Specific gene-expression analysis (**Figure 1D and 1E**) observed 25 differentially expressed genes (differential expression was defined as p = 0.02 and log2 fold change of 0.4, genes listed in **Supplemental Table S4**). As expected, genes associated with SARS-CoV-2 viral entry (*ACE2, FURIN*, and *S* – spike protein) and viral replication (*ORF1ab_REV*) were upregulated in the myocardium of patients with COVID-19. *DHX58* and *IL10RB*, both of which are associated with immune regulation, were also differentially expressed. *CT45A1* and *ID1* were the only genes noted to be downregulated with SARS-CoV-2 infection. Interestingly, the burden of SARS-CoV-2 expression (i.e. SARS-CoV-2 “High” vs “Low”) did not alter global myocardial expression patterns (**Supplemental Figure S3**). These results did not differ when data was analyzed per patient (**Supplemental Figure S4A**). Such observations highlight unique transcriptional programming within the myocardium of COVID-19 patients, independent of local viral load, and support the hypothesis that SARS-CoV-2 directly infects cardiac tissue.

### SARS-CoV-2 infection induces cell specific inflammation of coronary capillary endothelial cells

To further investigate myocardial transcriptional patterns, we examined regional differences based upon SARS-CoV-2 nucleocapsid expression among COVID-19 patients. Specifically, we leveraged the selective power of ROI segmentation to isolate areas where expression of the SARS-CoV-2 nucleocapsid was evident (labeled *positive*, **Figure 1F**). Compared to regions without SARS-CoV-2 nucleocapsid expression (i.e. *negative*), PCA of SARS-CoV-2 positive tissue revealed pattern clustering (**Figure 1G**) and differential gene expression patterns (**Figure 1H**). Interestingly, local SARS-CoV-2 infection upregulated mediators of viral antigen presentation including human leukocyte antigen (HLA) complex genes *HLA-B, HLA-I, HLA-DRB*, as well as the β_2_ microglobulin gene (*B2M*). Additionally, the immunomodulatory genes transmembrane protein 2 (*IFITM2*) and alpha-2 macroglobulin (*A2M*) were both upregulated in regions of viral infection. Infected cell types appeared to primarily be capillary endothelial cells as differentially expressed genes included the endothelial cell markers platelet endothelial cell adhesion molecule (*PECAM1 / CD31*) and endoglin (*ENG*). While these measurements did not specifically isolate transcripts from infected capillary endothelial cells, our combined genomic and anatomic assessment implicates coronary capillaries to be the predominant cellular structure infected by SARS-CoV-2 in the myocardium.

### Severe COVID-19 is not associated with differential programming of coronary arterial or venous endothelium

Based upon our finding that SARS-CoV-2 infection induces capillary specific inflammation, we attempted to isolate coronary vascular endothelial cells for gene expression analysis among the three patient groups. Unique from recent investigations of endothelial function during acute SARS-CoV-2 infection^8, 9, 13, 17^, our approach enabled us to investigate endothelial programming in native myocardial tissue without disruption of the surrounding anatomy or cellular microenvironment. Coronary arterial and venous endothelial cells were identified by anti-CD31 staining and isolated for spatial transcriptomics (**Figure 2A**). Among the 1860 genes tested, 1302 genes were above the limit of quantification. Despite marked heterogeneity in viral burden, PCA failed to show unique expression patterns between patient cohorts (**Figure 2B**). These findings were consistent across SARS-CoV-2 Low and High groups, and in keeping with our results of global myocardial profiling (**Supplemental Figure S3**). These results did not differ when data was analyzed per patient (**Supplemental Figure S4B**). Similarly, minimal differences in transcriptional programming were observed as only 9 endothelial genes were differentially expressed between Controls and all COVID-19 patients (**Figure 2C**). Of note, carcinoembryonic antigen-related cell adhesion molecule (*CEACAM3*) has been implicated in COVID-19 severity by regulating cell-cell communication of circulating neutrophils^18^, and was one of two upregulated endothelial genes measured during our study. Nonetheless, our findings highlight the coronary capillary bed, as opposed to arterial or venous systems, as potential mediators of myocardial injury and cardiovascular events associated with severe COVID-19.

**Figure 2.**
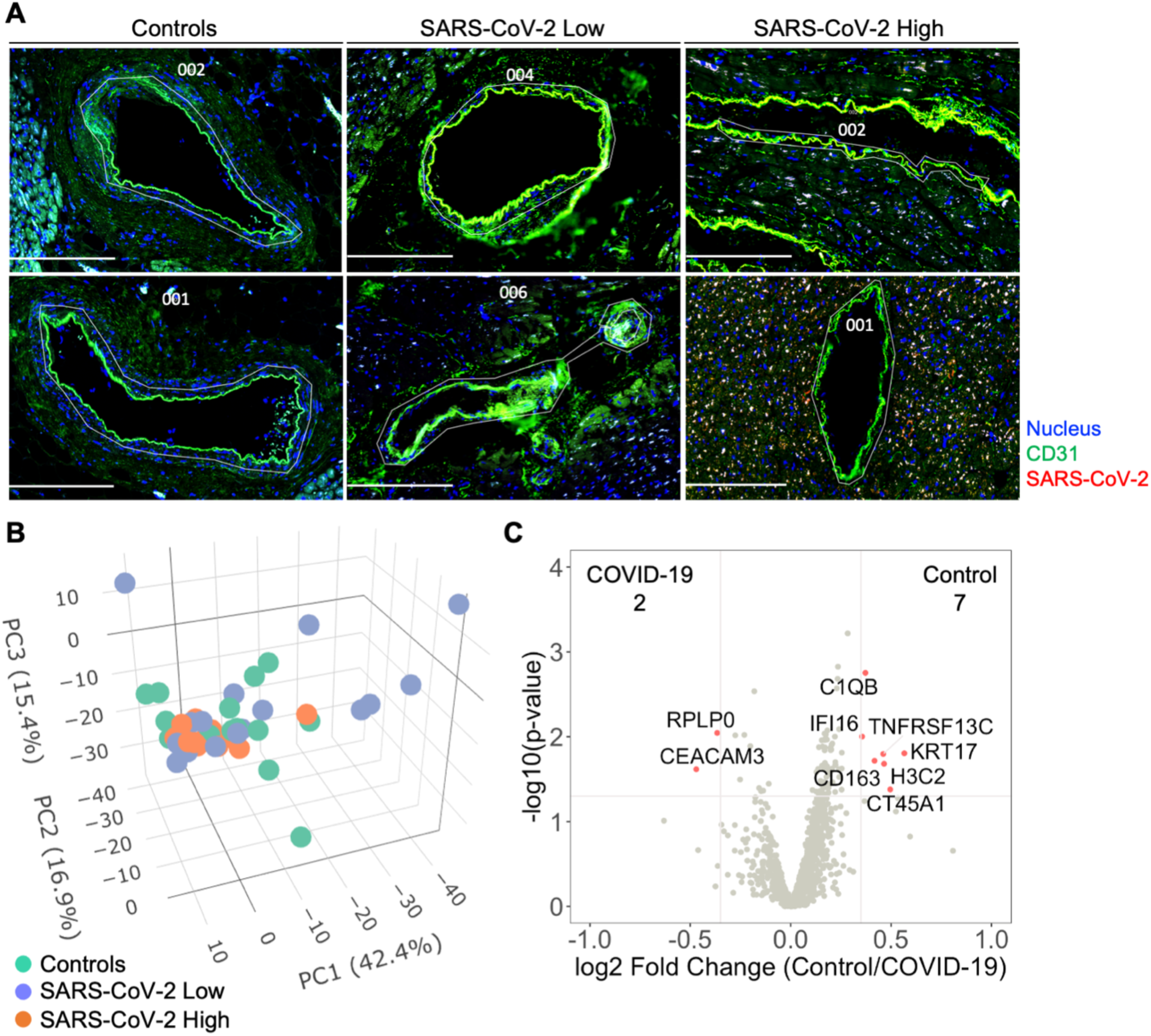
Severe COVID-19 is not associated with differential programming of the coronary arterial or venous endothelium. Coronary vascular endothelial cells were identified by immunofluorescence and selectively sampled for spatial transcriptomics (A, white custom tracings illustrate representative replicate ROIs). In addition to control samples, COVID-19 patient samples were segregated based upon relative expression of SARS-CoV-2 nucleocapsid. 1302 genes were above the limit of quantification. Despite marked heterogeneity in SARS-CoV-2 expression, principal component analysis failed to show pattern clustering (B). Similarly, minimal differences in transcriptional programming were observed as only 9 genes were observed to be differentially expressed between control and COVID-19 coronary endothelium (C, data includes ROIs from SARS-CoV-2 Low and High samples). Differential gene expression was defined as p = 0.02 and log2 fold change of 0.4. COVID-19 (n = 8 patients, 30 ROIs) and Control (n = 4 patients, 15 ROIs). Scale bars are 250µm for each image.

## DISCUSSION

Cardiovascular events are frequent complications associated with COVID-19^2, 7^. To date, investigations have centered on understanding the effect of SARS-CoV-2 on endothelial and myocardial function. While initial hypotheses suggested SARS-CoV-2 may induce a viral-mediated endotheliitis^3, 8^, recent evidence suggest otherwise^9^. Similarly, contrasting observations of direct SARS-CoV-2 infection of the myocardium have been reported. These discordant observations provided the rationale for our investigation seeking to characterize direct endothelial and/or myocardial mechanisms linking severe COVID-19 to reported cardiovascular events.

To our knowledge, this investigation is the first to measure transcriptional profiles of coronary endothelial cells from patients with COVID-19 using a novel spatial transcriptomics platform. As such, our findings represent the first *in situ* assessment of the transcriptome in coronary endothelial cells in the presence of SARS-CoV-2 virus. A critical observation from this work is the predominance of capillary endothelial infection and inflammation with SARS-CoV-2 as opposed to the endothelium of larger coronary vessels. Interestingly, diffuse endothelial activation was absent despite severe COVID-19 symptoms and confirmed viral infection of the surrounding tissue. As such, our work suggests that viral-mediated endotheliitis and endothelial activation is minimal and therefore unlikely to significantly modulate thromboembolic and cardiovascular events during COVID-19 infection. Other key observations from this work include 1) SARS-CoV-2 has variable expression patterns within the myocardium of COVID-19 patients, 2) viral infection induces a unique myocardial programming independent of local viral burden, and 3) myocardial tissues directly infected with SARS-CoV-2 have unique, pro-inflammatory expression profiles.

Similar to previous investigations^12^, results from our study did not observe classic hallmarks of myocarditis including inflammatory cell infiltrates on histologic review of COVID-19 patients (**Supplemental Figure S2**). Despite heterogeneous viral expression, spatial transcriptomic analysis identified unique transcriptional programming associated with all COVID-19 patients when compared to controls (**Figure 1C, 1D, and 1E**). While such changes have been previously reported within other patient cohorts^13^ and models of COVID-19 myocarditis^14^, our investigation presents the first such results from *in situ* tissue without technical manipulation. Our data show upregulation of genes within the myocardium associated with viral entry, viral replication, and immunomodulation. Notably, *IL10RB* encodes for interleukin (IL) - 10 receptor subunit B, which has been associated with poor clinical outcomes and suggested to be a critical regulator of host susceptibility to SARS-CoV-2^19^. The observed programming was independent of viral burden within the tissue, as all COVID-19 patients clustered together on principle component analysis (**Figure 1C**) and differential expression was not observed between SARS-CoV-2 High and Low patient cohorts (**Supplemental Figure S3**). Such observations suggest that SARS-CoV-2 may directly alter myocardial function, independent of classical hallmarks of myocarditis.

A unique aspect of our approach was the ability to selectively sample myocardial tissues with and without viral infection via segmentation. Interestingly, we observed cell specific differential programming that was influenced by SARS-CoV-2. As would be expected, SARS-CoV-2 infection upregulated mediators of viral antigen presentation including HLA haplotypes *HLA-B, HLA-I, HLA-DRB*. This observation is important as *HLA-B* and *HLA-DRB* have both been reported to influence COVID-19 disease severity^20, 21^ in association with preexisting medical comorbidities such as cardiovascular disease^22^. *B2M*, a component of MHC class 1 molecules, was also upregulated in infected cells and has been implicated as a risk marker for coronary heart disease and stroke^23^. Together, these findings expand our recognition of HLA alleles and MHC class 1 components that may play a relevant role in cardiovascular complications associated with SARS-CoV-2 infection.

Recently, our group identified altered coagulation pathways in pulmonary vessels from patients with severe COVID-19^16^. While differential programming of endothelial cells may have contributed to our observations, investigations from McCracken *et al*.^*9*^ and Johnson *et al*.^*10*^ failed to observe significant replicative infection or activation of human endothelial cells by SARS-CoV-2. Our present investigation of the coronary endothelium suggests an interesting anatomic paradigm between local capillaries and larger coronary vessels in response to COVID-19. While we did not observe significant changes in endothelial cell programming of coronary vessels (**Figure 2**), we did observe significant colocalization of the SARS-CoV-2 nucleocapsid and capillary endothelial cells suggesting potential viral infection (**Figure 1**). The endothelial genes *PECAM1 / CD31* and *ENG* were both enriched in regions of viral infection, further supporting our assertion of viral infection at the capillary bed (**Figure 1H**). Importantly, these infected endothelial cells showed upregulation of fibronectin-1 gene (*FN1*), which has been extensively linked with thrombus formation in injured arterioles^24^. Our results are aligned with prior investigations demonstrating increased ACE2 / TMPRSS2 expression in capillaries of COVID-19 patients^25^, and therefore support capillaritis as a prominent mediator of viral-induced myocardial injury. While these observations are notable, they fall short of supporting the original paradigm of large vessel endothelial activation as a mediator of cardiovascular events. Hence, despite upregulation of COVID-19 specific genes within the myocardium (*e*.*g. ACE2, S-*Spike, *Furin*) we still saw no differences in coronary endothelial cell programming to associate with clinical cardiovascular events. These results provide strong evidence that widespread endothelial activation is not a mediating component of cardiovascular events associated with COVID-19.

## STUDY LIMITATIONS

Our investigation has several limitations that are worth addressing. The most notable limitation is that all COVID-19 patients were likely infected with the alpha variant of SARS-CoV-2 as patient autopsies were conducted in 2020. While we speculate that our results are germane to more recent variants of SARS-CoV-2, our study was not designed to assess such differences. In addition, the minority of COVID-19 patients had documented thrombotic events and myocardial injury. While future studies could select more patients with cardiovascular events, the lack of observed differences in larger coronary vessels argues against the existence of meaningful biological differences.

Our observation of distinct capillary and arterial programming also warrants further discussion. Notably, pericytes adjacent to capillary endothelial cells have been reported to have relatively high ACE2 expression (compared to endothelial cells) and postulated to be a significant mediator for SARS-CoV-2 coagulopathy^9^. This confounder is difficult to eliminate utilizing the digital spatial platform due to limited morphology markers to differentiate pericytes and endothelial cells as well as the inability to isolate single cells for profiling. While our approach fails to definitively identify altered capillary endothelial reprogramming, the results nonetheless highlight the need for future studies to investigate the differences in coronary capillary response to SARS-CoV-2 infection.

Finally, our use of spatial transcriptomics limited the sample size of our investigation (*n* = 4 control and 8 COVID-19). While this is an important limitation, the clustering of groups based on principal component analysis (**Figures 1 and 2**) is reassuring that our analysis has not included significant outliers and may reflect true differences between control and COVID-19 patients. Future studies utilizing similar patient cohorts may help to extend our current observations.

## CONCLUSION

In summary, presented data identify unique regional and viral dependent differences within the myocardium and capillary bed that may influence the risk of cardiovascular events associated with SARS-CoV-2 infection. However, substantial differential expression patterns from the endothelium of larger coronary vessels were not observed. These results diminish endotheliitis and altered endothelial transcription as possible mechanisms for cardiovascular events associated with acute COVID-19.

## PERSPECTIVES

### Competency in medical knowledge

Identification and appropriate treatment of patients at greatest risk for severe cardiovascular complications is crucial to reduce mortality associated with acute COVID-19 infection. Hence, a comprehensive understanding of the vascular pathophysiology associated SARS-CoV-2 infection is paramount. The current investigation not only identifies novel coronary vascular molecular targets, but also describes coronary capillaritis as a hallmark of severe COVID-19. Furthermore, these findings highlight proinflammatory changes independent of local viral tissue expression.

### Translational outlook

A remaining obstacle to mitigating cardiovascular events associated with COVID-19 is our limited understanding of the mediators that influence SARS-CoV-2 infection of the myocardium and vasculature. We observed unique transcriptional programming associated with SARS-CoV-2 infection. Such differentially expressed genes warrant further mechanistic investigation to translate the current findings into potentially novel clinical targets. Similarly, alternative mechanisms of thromboembolic events (independent of endothelial activation) should garner increased attention given our observation of limited endothelial activation in response to SARS-CoV-2 infection. Separate from COVID-19, results from this work may have additional translational implications for other areas of cardiovascular research as it describes novel techniques to measure vascular transcriptional programming in patient samples without disruption of the surrounding microenvironment.

## List of Abbreviations (not counted in word count)

(SARS-CoV-2): Severe acute respiratory syndrome coronavirus-2
(ACE2): Angiotensin Converting Enzyme 2
(ROIs): regions of interests
(LOQ): limit of quantification
(PCA): principal component analysis
(HLA): human leukocyte antigen
(*B2M*): β_2_ microglobulin gene
(*IFITM2*): transmembrane protein 2
(*A2M*): alpha-2 macroglobulin
(*PECAM1 / CD31*): platelet endothelial cell adhesion molecule
(*ENG*): endoglin
(*CEACAM3*): carcinoembryonic antigen-related cell adhesion molecule
(*FN1*): fibronectin-1 gene

## ACKNOWLEDGMENTS

We would like to thank the Tissue Biorepository Core Facility at UAB for their assistance with the processing of patient specimens. Specifically, we thank Dezhi Wang for her assistance with tissue processing. Research reported in this publication was also supported by the UAB High Resolution Imaging Facility. Specifically, we thank Shawn Williams for his assistance with initial confocal imaging. We also thank Drs. Nathan Erdmann, Kevin Harrod, and J. Edwin Blalock for their insightful feedback regarding this manuscript. Lastly, we thank Beth Thompson for administrative support.

## SUPPLEMENTAL MATERIALS

Supplemental Methods

Supplemental Table S1, S2, S3, S4

Supplemental Figures S1, S2, S3, S4

Data Set

Major Resources Table

## REFERENCES

1. Wiersinga WJ, Rhodes A, Cheng AC, Peacock SJ, Prescott HC. Pathophysiology, transmission, diagnosis, and treatment of coronavirus disease 2019 (covid-19): A review. JAMA. 2020;324:782–793

2. Nishiga M, Wang DW, Han Y, Lewis DB, Wu JC. Covid-19 and cardiovascular disease: From basic mechanisms to clinical perspectives. Nat Rev Cardiol. 2020;17:543–558

3. Varga Z, Flammer AJ, Steiger P, Haberecker M, Andermatt R, Zinkernagel AS, Mehra MR, Schuepbach RA, Ruschitzka F, Moch H. Endothelial cell infection and endotheliitis in covid-19. Lancet. 2020;395:1417–1418

4. McFadyen JD, Stevens H, Peter K. The emerging threat of (micro)thrombosis in covid-19 and its therapeutic implications. Circ Res. 2020;127:571–587

5. Teuwen LA, Geldhof V, Pasut A, Carmeliet P. Covid-19: The vasculature unleashed. Nat Rev Immunol. 2020;20:389–391

6. Watanabe T, Barker TA, Berk BC. Angiotensin ii and the endothelium: Diverse signals and effects. Hypertension. 2005;45:163–169

7. Liu PP, Blet A, Smyth D, Li H. The science underlying covid-19: Implications for the cardiovascular system. Circulation. 2020;142:68–78

8. Liu F, Han K, Blair R, Kenst K, Qin Z, Upcin B, Worsdorfer P, Midkiff CC, Mudd J, Belyaeva E, Milligan NS, Rorison TD, Wagner N, Bodem J, Dolken L, Aktas BH, Vander Heide RS, Yin XM, Kolls JK, Roy CJ, Rappaport J, Ergun S, Qin X. Sars-cov-2 infects endothelial cells in vivo and in vitro. Front Cell Infect Microbiol. 2021;11:701278

9. McCracken IR, Saginc G, He L, Huseynov A, Daniels A, Fletcher S, Peghaire C, Kalna V, Andaloussi-Mae M, Muhl L, Craig NM, Griffiths SJ, Haas JG, Tait-Burkard C, Lendahl U, Birdsey GM, Betsholtz C, Noseda M, Baker AH, Randi AM. Lack of evidence of angiotensin-converting enzyme 2 expression and replicative infection by sars-cov-2 in human endothelial cells. Circulation. 2021;143:865–868

10. Johnson JE, McGuone D, Xu ML, Jane-Wit D, Mitchell RN, Libby P, Pober JS. Coronavirus disease 2019 (covid-19) coronary vascular thrombosis: Correlation with neutrophil but not endothelial activation. Am J Pathol. 2022;192:112–120

11. Omidi F, Hajikhani B, Kazemi SN, Tajbakhsh A, Riazi S, Mirsaeidi M, Ansari A, Ghanbari Boroujeni M, Khalili F, Hadadi S, Nasiri MJ. Covid-19 and cardiomyopathy: A systematic review. Front Cardiovasc Med. 2021;8:695206

12. Delorey TM, Ziegler CGK, Heimberg G, Normand R, Yang Y, Segerstolpe A, Abbondanza D, Fleming SJ, Subramanian A, Montoro DT, Jagadeesh KA, Dey KK, Sen P, Slyper M, Pita-Juarez YH, Phillips D, Biermann J, Bloom-Ackermann Z, Barkas N, Ganna A, Gomez J, Melms JC, Katsyv I, Normandin E, Naderi P, Popov YV, Raju SS, Niezen S, Tsai LT, Siddle KJ, Sud M, Tran VM, Vellarikkal SK, Wang Y, Amir-Zilberstein L, Atri DS, Beechem J, Brook OR, Chen J, Divakar P, Dorceus P, Engreitz JM, Essene A, Fitzgerald DM, Fropf R, Gazal S, Gould J, Grzyb J, Harvey T, Hecht J, Hether T, Jane-Valbuena J, Leney-Greene M, Ma H, McCabe C, McLoughlin DE, Miller EM, Muus C, Niemi M, Padera R, Pan L, Pant D, Pe’er C, Pfiffner-Borges J, Pinto CJ, Plaisted J, Reeves J, Ross M, Rudy M, Rueckert EH, Siciliano M, Sturm A, Todres E, Waghray A, Warren S, Zhang S, Zollinger DR, Cosimi L, Gupta RM, Hacohen N, Hibshoosh H, Hide W, Price AL, Rajagopal J, Tata PR, Riedel S, Szabo G, Tickle TL, Ellinor PT, Hung D, Sabeti PC, Novak R, Rogers R, Ingber DE, Jiang ZG, Juric D, Babadi M, Farhi SL, Izar B, Stone JR, Vlachos IS, Solomon IH, Ashenberg O, Porter CBM, Li B, Shalek AK, Villani AC, Rozenblatt-Rosen O, Regev A. Covid-19 tissue atlases reveal sars-cov-2 pathology and cellular targets. Nature. 2021;595:107–113

13. Brauninger H, Stoffers B, Fitzek ADE, Meissner K, Aleshcheva G, Schweizer M, Weimann J, Rotter B, Warnke S, Edler C, Braun F, Roedl K, Scherschel K, Escher F, Kluge S, Huber TB, Ondruschka B, Schultheiss HP, Kirchhof P, Blankenberg S, Puschel K, Westermann D, Lindner D. Cardiac sars-cov-2 infection is associated with pro-inflammatory transcriptomic alterations within the heart. Cardiovasc Res. 2021

14. Bailey AL, Dmytrenko O, Greenberg L, Bredemeyer AL, Ma P, Liu J, Penna V, Winkler ES, Sviben S, Brooks E, Nair AP, Heck KA, Rali AS, Simpson L, Saririan M, Hobohm D, Stump WT, Fitzpatrick JA, Xie X, Zhang X, Shi PY, Hinson JT, Gi WT, Schmidt C, Leuschner F, Lin CY, Diamond MS, Greenberg MJ, Lavine KJ. Sars-cov-2 infects human engineered heart tissues and models covid-19 myocarditis. JACC Basic Transl Sci. 2021;6:331–345

15. Phillips N. The coronavirus is here to stay - here’s what that means. Nature. 2021;590:382–384

16. Margaroli C, Benson P, Sharma NS, Madison MC, Robison SW, Arora N, Ton K, Liang Y, Zhang L, Patel RP, Gaggar A. Spatial mapping of sars-cov-2 and h1n1 lung injury identifies differential transcriptional signatures. Cell Rep Med. 2021;2:100242

17. Guervilly C, Burtey S, Sabatier F, Cauchois R, Lano G, Abdili E, Daviet F, Arnaud L, Brunet P, Hraiech S, Jourde-Chiche N, Koubi M, Lacroix R, Pietri L, Berda Y, Robert T, Degioanni C, Velier M, Papazian L, Kaplanski G, Dignat-George F. Circulating endothelial cells as a marker of endothelial injury in severe covid -19. J Infect Dis. 2020;222:1789–1793

18. Huang R, Meng T, Zha Q, Cheng K, Zhou X, Zheng J, Zhang D, Liu R. The predicting roles of carcinoembryonic antigen and its underlying mechanism in the progression of coronavirus disease 2019. Crit Care. 2021;25:234

19. Voloudakis G, Hoffman G, Venkatesh S, Lee KM, Dobrindt K, Vicari JM, Zhang W, Beckmann ND, Jiang S, Hoagland D, Bian J, Gao L, Corvelo A, Cho K, Lee JS, Iyengar SK, Luoh SW, Akbarian S, Striker R, Assimes TL, Schadt EE, Merad M, tenOever BR, Charney AW, Mount Sinai C-B, Initiative VAMVPC-S, Brennand KJ, Lynch JA, Fullard JF, Roussos P. Il10rb as a key regulator of covid-19 host susceptibility and severity. medRxiv. 2021

20. Langton DJ, Bourke SC, Lie BA, Reiff G, Natu S, Darlay R, Burn J, Echevarria C. The influence of hla genotype on the severity of covid-19 infection. HLA. 2021;98:14–22

21. Weiner J, Suwalski P, Holtgrewe M, Rakitko A, Thibeault C, Muller M, Patriki D, Quedenau C, Kruger U, Ilinsky V, Popov I, Balnis J, Jaitovich A, Helbig ET, Lippert LJ, Stubbemann P, Real LM, Macias J, Pineda JA, Fernandez-Fuertes M, Wang X, Karadeniz Z, Saccomanno J, Doehn JM, Hubner RH, Hinzmann B, Salvo M, Blueher A, Siemann S, Jurisic S, Beer JH, Rutishauser J, Wiggli B, Schmid H, Danninger K, Binder R, Corman VM, Muhlemann B, Arjun Arkal R, Fragiadakis GK, Mick E, Comet C, Calfee CS, Erle DJ, Hendrickson CM, Kangelaris KN, Krummel MF, Woodruff PG, Langelier CR, Venkataramani U, Garcia F, Zyla J, Drosten C, Alice B, Jones TC, Suttorp N, Witzenrath M, Hippenstiel S, Zemojtel T, Skurk C, Poller W, Borodina T, Pa-Covid SG, Ripke S, Sander LE, Beule D, Landmesser U, Guettouche T, Kurth F, Heidecker B. Increased risk of severe clinical course of covid-19 in carriers of hla-c*04:01. EClinicalMedicine. 2021;40:101099

22. Mobini Kesheh M, Shavandi S, Hosseini P, Kakavand-Ghalehnoei R, Keyvani H. Bioinformatic hla studies in the context of sars-cov-2 pandemic and review on association of hla alleles with preexisting medical conditions. Biomed Res Int. 2021;2021:6693909

23. Prentice RL, Zhao S, Johnson M, Aragaki A, Hsia J, Jackson RD, Rossouw JE, Manson JE, Hanash SM. Proteomic risk markers for coronary heart disease and stroke: Validation and mediation of randomized trial hormone therapy effects on these diseases. Genome Med. 2013;5:112

24. Ni H, Yuen PS, Papalia JM, Trevithick JE, Sakai T, Fassler R, Hynes RO, Wagner DD. Plasma fibronectin promotes thrombus growth and stability in injured arterioles. Proc Natl Acad Sci U S A. 2003;100:2415–2419

25. Maccio U, Zinkernagel AS, Shambat SM, Zeng X, Cathomas G, Ruschitzka F, Schuepbach RA, Moch H, Varga Z. Sars-cov-2 leads to a small vessel endotheliitis in the heart. EBioMedicine. 2021;63:103182

