## Supplemental Document for "Spatial transcriptomic profiling of coronary endothelial cells in SARS-CoV-2 myocarditis"

**SOURCES OF FUNDING:** CM was supported by the Cystic Fibrosis Foundation – Postdoc-to-Faculty Award (MARGAR21F5), GAP was supported by the American Heart Association – Amos Medical Faculty Development Program (AHA-AMFDP) award 18AMFDP34380568, National Heart Lung and Blood Institute (NHLBI) 3R35HL135710, and the UAB Comprehensive Cardiovascular Center. Experimental work was supported by the AHA-AMFDP (18AMFDP34380568) and NHLBI (R01HL102371). This work was also supported by the Veterans Affairs Medical Center Grant I01BX001756-02 (to AG).

**RELATIONSHIP TO INDUSTRY AND DISCLOSURES:** The authors declare no conflict of interest

**ADDRESS FOR CORRESPONDENCE:**
**Gregory A. Payne MD PhD**
Zeigler Research Building 531
1720 2nd Avenue South
Birmingham, AL 35294

### SUPPLEMENTAL METHODS

**Materials and data availability.** Anonymized datasets generated during this study will be available on Mendeley Data. Other requests for resources and reagents are available from the corresponding author upon reasonable request. Requests should be directed to and will be fulfilled by Dr. Gregory Payne. This study did not generate new or unique reagents. Please see the **Major Resources Table** in the Supplemental Materials for detailed experimental reagents.

**Human subjects.** The study was approved by an institutional review board (UAB-IRB 300005258, VA-IRB 1573682) and summary demographic and clinical data are presented in **Tables S1 and S2**. The autopsy authorization from next of kin included consent for use of the tissue for research purposes. SARS-CoV-2 autopsy samples were collected from patients that died due to pneumonia and acute respiratory failure between March and June of 2020. All SARS-CoV-2 participants had chest imaging consistent with findings of pneumonia and positive nucleic acid amplification test (PCR) for SARS-CoV-2 indicating COVID-19 pneumonia. Myocardial specimens were stained for SARS-CoV-2 nucleocapsid and stratified into quartiles based upon relative mean fluorescent intensities (MFI) measured by a Nikon A1R confocal microscope. Specimen in the bottom quartile (<25% MFI) were defined as SARS-CoV-2 "Low", while the top quartile (>75% MFI) was defined as SARS-CoV-2 "high". Separately, four deceased patients without infection or known cardiovascular disease who died before the existence of SARS-CoV-2 were included as control subjects (all died prior to 2020). All participants were grouped according to COVID-19 status and myocardial SARS-CoV-2 expression as discussed.

**Histology and Immunofluorescence.** The presence of myocardial infection with SARS-CoV-2 was determined prior to GeoMX digital spatial profiling by immunofluorescence to the viral nucleocapsid. Specifically, all myocardial tissue specimens for IHC staining were fixed with 10% neutral buffered formalin (Fisher Scientific) at room temperature for 24 hours. Samples were then dehydrated, and paraffin embedded prior to serial 5 µm thick sectioning. Sections were dried overnight at 60°C, deparaffinized and hydrated using graded concentrations of ethanol to deionized water. Tissue sections for IHC were subjected to antigen retrieval by 0.01 M Tris-1 mM EDTA buffer (pH 9) in steamer for 5 minutes (buffer preheated with the steam setting for 20 minutes). Following antigen retrieval, all sections were washed gently in deionized water. Sections were soaked in PBS for at least 10 minutes. 3% BSA was then applied on sections for 40 min at RT to block non-specific binding followed by "blot dry" (no rinse after blocking at this step). Anti-SARS-CoV-2 nucleocapsid (GeneTex, GTX135361; 1:500) and anti-CD31 (Abcam, ab9498, 1:200) antibodies were directly conjugated to fluorescent markers (see ***GeoMX digital spatial profiling*** below) and diluted in 3% BSA prior to application in a dark incubation chamber for one hour at RT. Sections were then rinsed with PBS 5 minutes, 3 times with agitation and then blotted dry. Nuclei were counter stained with DAPI (Biolegend, stock 100ng/ml and 1:1000 in PBS working solution) for 15 minutes in the dark. Finally, sections were washed with PBS prior to addition of mounting media with DAPI (ThermoFisher, ProLong Gold antifade) and coverslip. Confocal immunofluorescence images were acquired using the Nikon A1R confocal microscope. Myocardial tissue from patients without SARS-CoV-2 were used as negative controls for Anti-SARS-CoV-2 nucleocapsid and showed no appreciable signal.

**GeoMX digital spatial profiling. Figure S1** illustrates the work-flow utilized for digital spatial profiling of myocardial tissues. Briefly, paraffin embedded tissues were processed and analyzed locally using a combination of fluorescently labeled anti-SARS-CoV-2 nucleocapsid (GeneTex, GTX135361; 1:500) and anti-CD31 (Abcam, ab9498, 1:200) antibodies. Anti-SARS-CoV-2 was conjugated to PE / R-Phycoerythrin (Expedeon Lightning-Link R-PE Conjugation Kit / Abcam, ab102918), while anti-CD31 was linked to Alexa Fluor 488 (Abcam, Lighting-Link ab236553). Nuclei were counterstained with SYTO61 (ThermoFisher, S11343, 1:1000 in PBS working solution). Fluorescent antibodies were combined with the GeoMX Cancer Transcriptome (Nanostring) and COVID-19 Immune Response Atlas gene sets with custom probes specific for SARS-CoV-2 lung infection and tissue responses (see **Table S3** for SARS-CoV-2 related gene list), totaling 1860 genes. Selection of regions of interests (ROIs) was performed based on 1) immunofluorescent viral staining, 2) cellular immunofluorescent profile and anatomic features consistent with coronary vascular tissue, and 3) cellular anatomic features consistent with myocardial tissue observed in the hematoxylin and eosin (H&E) stained sections. To ensure even and representative selection of ROIs, general myocardial regions were selected evenly across the entire tissue section. Endothelial ROIs were sampled whenever possible due to the heterogeneity of coronary arterial and venous vessels. **Figure S2** illustrates the distribution of all ROIs for each COVID-19 patient. For endothelial cell-specific profiling (**Figure 2**), at least 50 cells per ROI were utilized for analyses.

**Quantification.** Libraries of the oligo tags collected with the Nanostring GeoMx platform were prepared using the GeoMx Seq code plates and reagents (Nanostring Technologies) with unique I7 and I5 indices. Paired sequencing was performed on the NovaSeq 6000 (Flow cell S100, Medgenome) of the oligo tags and not on the transcript itself, providing a more accurate transcript count with less sequencing bias. FASTQ files were converted into DCC files using the online Nanostring pipeline available on Basespace (Illumina). RNA probe counts used in the analyses were selected following a sequencing QC according to Nanostring protocols. Specifically, counts from each identified region of interest are analyzed, and under-sequenced samples are dropped (field of view percentage of 75% and Binding density from 0.1 to 2.25). A QC probe was also used where mRNAs are targeted by multiple probes and outlier probes are dropped from downstream data analysis (positive spike-in normalization factory between 0.3 and 3) [1]. Then RNA counts were normalized using a signal-based normalization, in which individual counts are normalized against the 75$^{th}$ percentile of signal from their own region of interest. The final list of detectable genes was then obtained by dropping genes in each specific group (i.e. endothelial, COVID status, SARS-CoV-2 nucleocapsid expression) by using a limit of quantification (LOQ) of 20% coverage within replicates. The LOQ was calculated using the geometric mean and geometric standard deviation of negative probes in the dataset. Counts were normalized to log2 and statistical comparisons were performed using a mixed linear model with Benjamini-Hochberg correction to account for false discovery rate[1, 2].

**Statistical Analysis**. Clinical data is expressed as mean ± standard deviation or n (%) unless otherwise indicated. 1-way ANOVA or Chi square test used to measure differences between groups for continuous and categorical values, respectively. For digital spatial profiling, comparison of all groups (e.g., Control, SARS-CoV-2 Low, and SARS-CoV-2 High) was performed by comparing the technical replicates from each biological group. This approach was taken given the heterogeneous vascularity and presence of endothelial ROIs available per myocardial tissue section. Each figure illustrates replicate ROIs for a given patient sample size. Similar statistical analyses were performed per patient (n = 4 patients per group) without any difference to the final conclusions (see results below). Counts were normalized to log2 and statistical comparisons were performed using a mixed linear model with Benjamini-Hochberg correction to account for false discovery rate. P value threshold for differential gene expression was set at p = 0.02 and log2 fold change of 0.4. All details for the statistical analyses and number of replicates can be found in the figure legends. All analyses for the volcano plots can be found in the supplemental Data Set file.

**Independent data access and analysis:** Drs. Camilla Margaroli and Gregory Payne had full access to all the data in the study and take responsibility for its integrity and analysis.

**Table S1.** Clinical Characteristics by Group

| | Total<br><br>(n=12) | Controls<br><br>(n=4) | SARS-CoV-2 "Low" (n=4) | SARS-CoV-2 "High" (n=4) | p-value |
|---|---|---|---|---|---|
| Age, years | 67±13 | 62±8 | 66±10 | 75±17 | 0.34 |
| Male sex | 9 (75%) | 3 (75%) | 3 (75%) | 3 (75%) | 0.99 |
| Race<br>Caucasian<br>African American | <br>7 (58%)<br>5 (42$) | <br>2 (50%)<br>2 (50%) | <br>2 (50%)<br>2 (50%) | <br>3 (75%)<br>1 (25%) | <br>0.47 |
| Composite Cardiovascular Disease Diagnosis$ | 8 (67%) | 2 (50%) | 3 (75%) | 3 (75%) | 0.69 |
| Coronary artery disease | 3 (25%) | 1 (25%) | 0 (0%) | 2 (50%) | 0.26 |
| CHF | 4 (33%) | 2 (50%) | 0 (0%) | 2 (50%) | 0.22 |
| Hypertension | 7 (58%) | 2 (50%) | 3 (75%) | 2 (50%) | 0.71 |
| Diabetes | 3 (25%) | 1 (25%) | 2 (50%) | 0 (0%) | 0.26 |
| Lung disease | 4 (33%) | 0 (0%) | 2 (50%) | 2 (50%) | 0.22 |
| CKD or ESRD | 3 (25%) | 1 (25%) | 0 (0%) | 2 (50%) | 0.26 |
| Immunocompromised status | 2 (17%) | 0 (0%) | 1 (25%) | 1 (25%) | 0.55 |
| Mean arterial pressure, mmHg | 85±19 | 85±33 | 88±12 | 82±8 | 0.94 |
| Pulse, beats/min | 88±19 | 88±13 | 90±33 | 86±11 | 0.78 |
| Resp rate, breath/min | 23±5 | 22±6 | 25±5 | 24±3 | 0.94 |
| BNP / proBNP (pg/mL) | 282±154 | 239±89 | 305±229 | 299# | 0.93 |
| Troponin (ng/L) | 3264±5450 | 8222±11487 | 2581±4560 | 1469±1741 | 0.38 |
| LDH (U/L) | 598±462 | 1559# | 498±265 | 377±143 | 0.02 |
| CRP (mg/L) | 256±171 | 3.4# | 377±7.9 | 260±162 | 0.22 |
| Lactic Acid (mmol/L) | 5.5±4.3 | 8.1±6.1 | 4.8±2.4 | 3.0±1.6 | 0.30 |
| WBC ($10^3$ cells/L) | 11±4 | 12±6 | 11±2 | 10±4 | 0.89 |
| P/F ratio | 210±194 | 302±180 | 229±274 | 101±51 | 0.67 |
| A-a gradient | 278±231 | 127±150 | 307±247 | 398±248 | 0.26 |
| APACHE II score | 19±10 | 25±10 | 14±12 | 19±5 | 0.28 |
| Systemic glucocorticoid use | 6 (50%) | 1 (25%) | 3 (75%) | 2 (50%) | 0.37 |
| Remdesivir use | 1 (8%) | 0 (0%) | 1 (25%) | 0 (0%) | 0.34 |

Data expressed as mean ± standard deviation or n (%) unless otherwise indicated. 1-way ANOVA or Chi square test used to measure differences between groups for continuous and categorical values, respectively. $Composite cardiovascular disease diagnosis was a "yes" answer to any of the following: coronary artery disease, atherosclerosis, previous PCI, previous CABG, congestive heart failure, arrhythmia, cerebrovascular disease, peripheral arterial disease. #Indicates one data point in the group.

**Table S2.** Sequelae and Complications from Critical Illness.

| | Total<br><br>(n=12) | Control<br><br>(n=4) | SARS-CoV-2<br>"Low"<br>(n=4) | SARS-CoV-2<br>"High"<br>(n=4) | p-value |
|---|---|---|---|---|---|
| Atrial arrhythmia | 6 (50%) | 1 (25%) | 2 (50%) | 3 (75%) | 0.37 |
| Myocardial infarction | 3 (25%) | 1 (25%) | 1 (25%) | 1 (25%) | 0.99 |
| Ischemic stroke | 1 (8%) | 1 (25%) | 0 (0%) | 0 (0%) | 0.34 |
| Pulmonary embolism | 1 (8%) | 0 (0%) | 1 (25%) | 0 (0%) | 0.34 |
| Acute kidney injury | 7 (58%) | 3 (75%) | 2 (50%) | 2 (50%) | 0.71 |
| Abnormal LVEF on Echo | 2/9 (22%) | 1/3 (33%) | 0/4 (0%) | 1/2 (50%) | 0.33 |
| Abnormal RV on Echo | 4/9 (44%) | 1/3 (33%) | 3/4 (75%) | 0/2 (0%) | 0.20 |

Data expressed as mean ± standard deviation or n (%) unless otherwise indicated. 1-way ANOVA or Chi square test used to measure differences between groups for continuous and categorical values, respectively.

**Table S3. COVID-19 Specific Gene Set**

| | |
|---|---|
| S | Spike Protein |
| ORF1ab | ORF1ab |
| ORF1ab_REV | Negative Strand ORF1ab |
| ACE2 | angiotensin I converting enzyme 2 |
| ABCA3 | ATP binding cassette subfamily A member 3 |
| AQP5 | aquaporin 5 |
| DHX58 | DEXH (Asp-Glu-X-His) box polypeptide 58 |
| FURIN | furin (paired basic amino acid cleaving enzyme) |
| HAS2 | hyaluronan synthase 2 |
| HOPX | HOP homeobox |
| IFNLR1 | interferon lambda receptor 1 |
| IL10RB | interleukin 10 receptor subunit beta |
| MUC13 | mucin 13, cell surface associated |
| MUC2 | mucin 2, oligomeric mucus/gel-forming |
| MUC5AC | mucin 5AC, oligomeric mucus/gel-forming |
| MUC5B | mucin 5B, oligomeric mucus/gel-forming |
| NAPSA | napsin A aspartic peptidase |
| PGC | progastricsin (pepsinogen C) |
| SCGB1A1 | secretoglobin family 1A member 1 |
| SFTPA1 | surfactant protein A1 |
| SFTPB | surfactant protein B |
| SFTPC | surfactant protein C |
| SFTPD | surfactant protein D |
| TP63 | tumor protein p63 |
| CC2D1B | coiled-coil and C2 domain containing 1B |
| SF3A3 | splicing factor 3a subunit 3 |

A custom SARS-CoV-2 gene set was combined with the Cancer Transcriptome and Immune Response Atlas gene sets for GeoMX digital spatial profiling.

**Table S4. Myocardial Differentially Expressed Genes from COVID-19 Myocardium**

| | |
|---|---|
| ABCA3 | ATP binding cassette subfamily A member 3 |
| NAPSA | napsin A aspartic peptidase |
| IL10RB | interleukin 10 receptor subunit beta |
| SFTPC | surfactant protein C |
| SFTPA1 | surfactant protein A1 |
| ORF1ab_REV | Negative Strand ORF1ab |
| MUC13 | mucin 13, cell surface associated |
| SFTPD | surfactant protein D |
| MUC5AC | mucin 5AC, oligomeric mucus/gel-forming |
| FURIN | furin (paired basic amino acid cleaving enzyme) |
| TP63 | tumor protein p63 |
| PGC | progastricsin (pepsinogen C) |
| DHX58 | DEXH (Asp-Glu-X-His) box polypeptide 58 |
| MUC2 | mucin 2, oligomeric mucus/gel-forming |
| ACE2 | angiotensin I converting enzyme 2 |
| MUC5B | mucin 5B, oligomeric mucus/gel-forming |
| AQP5 | aquaporin 5 |
| S | Spike Protein |
| SCGB1A1 | secretoglobin family 1A member 1 |
| HOPX | HOP homeobox |
| CC2D1B | coiled-coil and C2 domain containing 1B |
| SFTPB | surfactant protein B |
| SF3A3 | splicing factor 3a subunit 3 |
| CT45A1 | cancer / Testis Antigen Family 45 Member A1 |
| ID1 | Inhibitor of DNA binding 1, HLH protein |

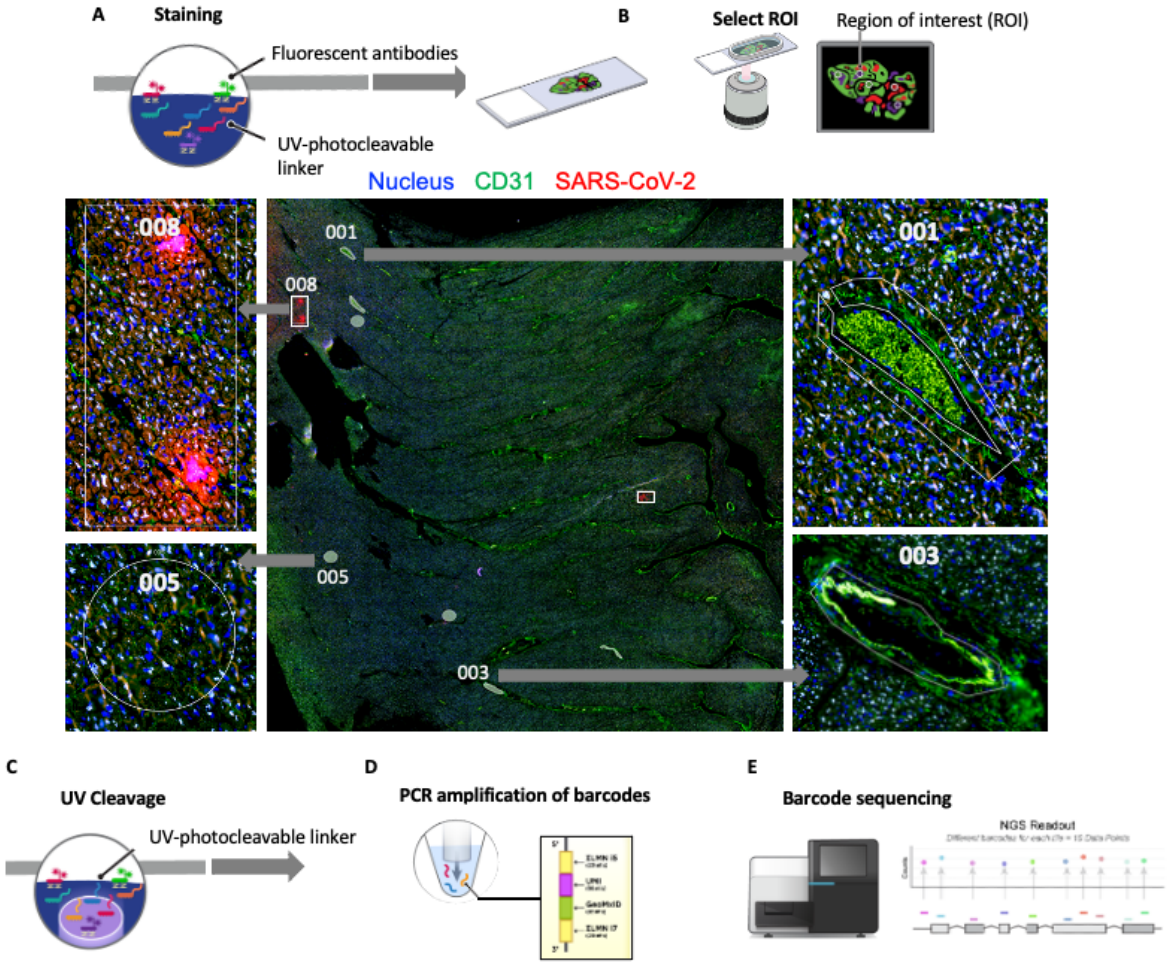


**Figure S1.** GeoMx Nanostring workflow. Myocardial autopsy samples were processed and stained using standard immunohistochemical protocol with both fluorescent labeled morphology markers and sequence specific RNA probes (A). Morphology markers (CD31 and SARS-CoV-2 nucleocapsid) were used to identify coronary endothelium and areas of SARS-CoV-2 expression as Regions of Interest (ROI). A representative myocardial section with selected ROIs is illustrated (B). After ROI selection, UV light is systematically shone on ROIs to photocleave and decouple barcodes from the RNA probes only from the selected area of interest (C). RNA probes for the ROI were aspirated and deposited into unique well of a 96 well plate (D). Five probes per mRNA target were used and were coupled to unique molecular barcodes to identify specific gene targets. The Unique Molecular Identifier (UMI) identifies a specific molecule and accounts for amplification bias from PCR. The PCR step adds dual indexing barcodes to identify ROIs and adds Illumina flow cell adapter regions to enable sequencing. The barcodes are sequenced and prepared for further analysis (E).

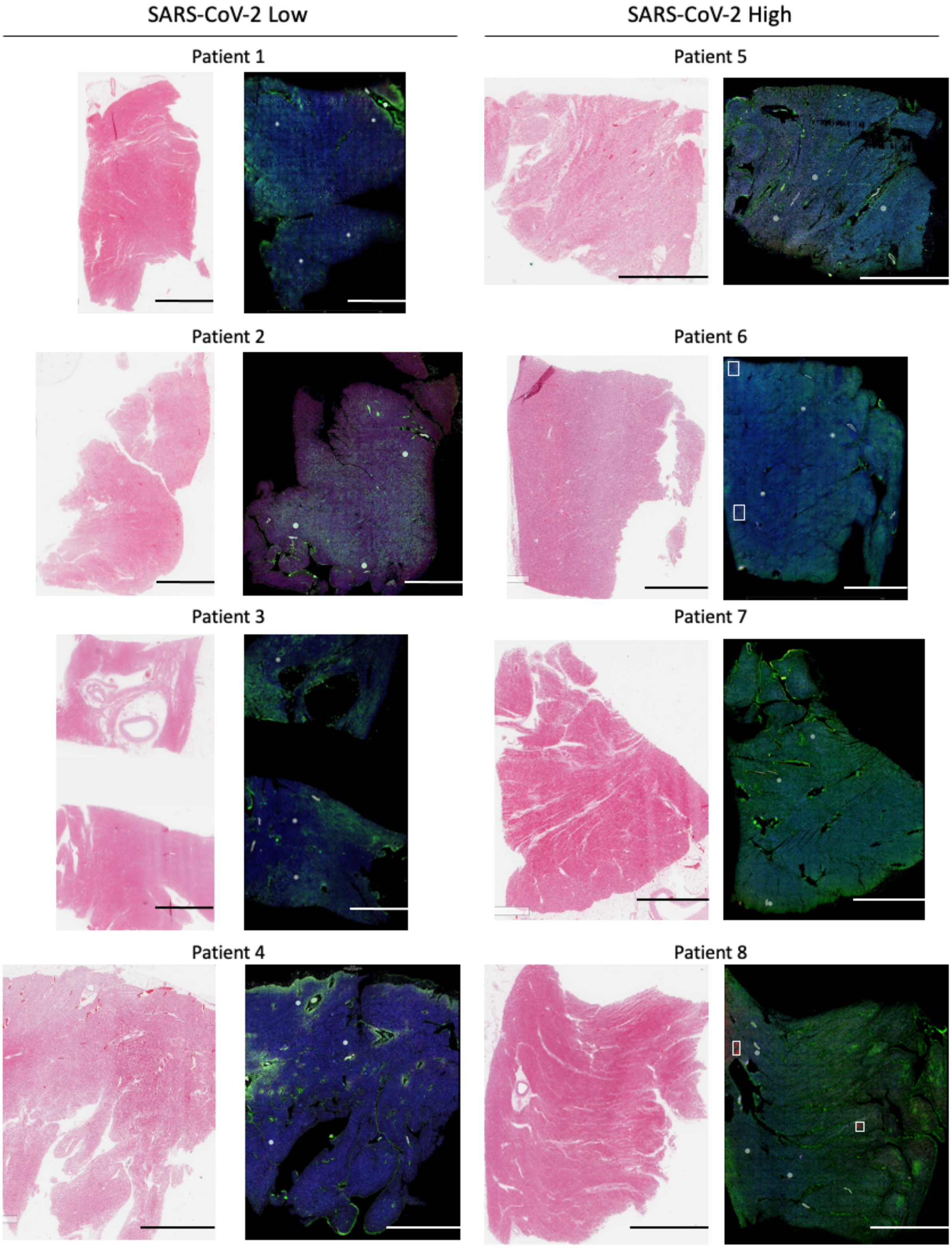


**Figure S2.** COVID-19 myocardial tissue sections and ROI selection. Representative H&E and fluorescently labeled tissue sections embedded in paraffin from all COVID-19 patients are illustrated. ROIs for each patient sample are highlighted by circles, rectangles, or custom shapes (used to identify coronary endothelial cells). Histologic review by a clinical anatomic pathologist failed to observe inflammatory cell infiltration in all COVID-19 tissues. Heterogeneous expression of SARS-CoV-2 was observed. Scale bar is 5mm for each image.

### MYOCARDIAL ROIs

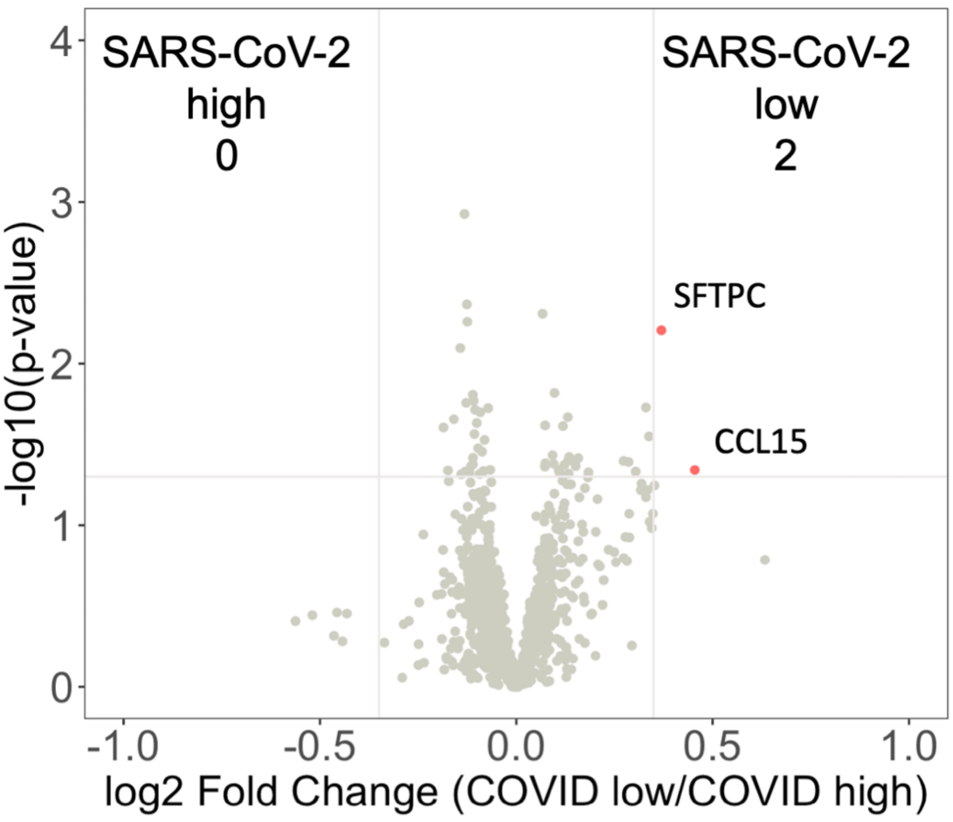


**Figure S3.** Intensity of SARS-CoV-2 expression does not alter general myocardial expression in severe COVID-19 infection. COVID-19 patient samples were segregated based upon relative expression of SARS-CoV-2 nucleocapsid. 1302 genes were above the limit of quantification. Minimal differences in transcriptional programming were observed between SARS-CoV-2 high and SARS-CoV-2 low patient samples. Differential gene expression was defined as p = 0.02 and log2 fold change of 0.4. SARS-CoV-2 high (n = 4 patients, 12 ROIs) and SARS-CoV-2 low (n = 4 patients, 12 ROIs).

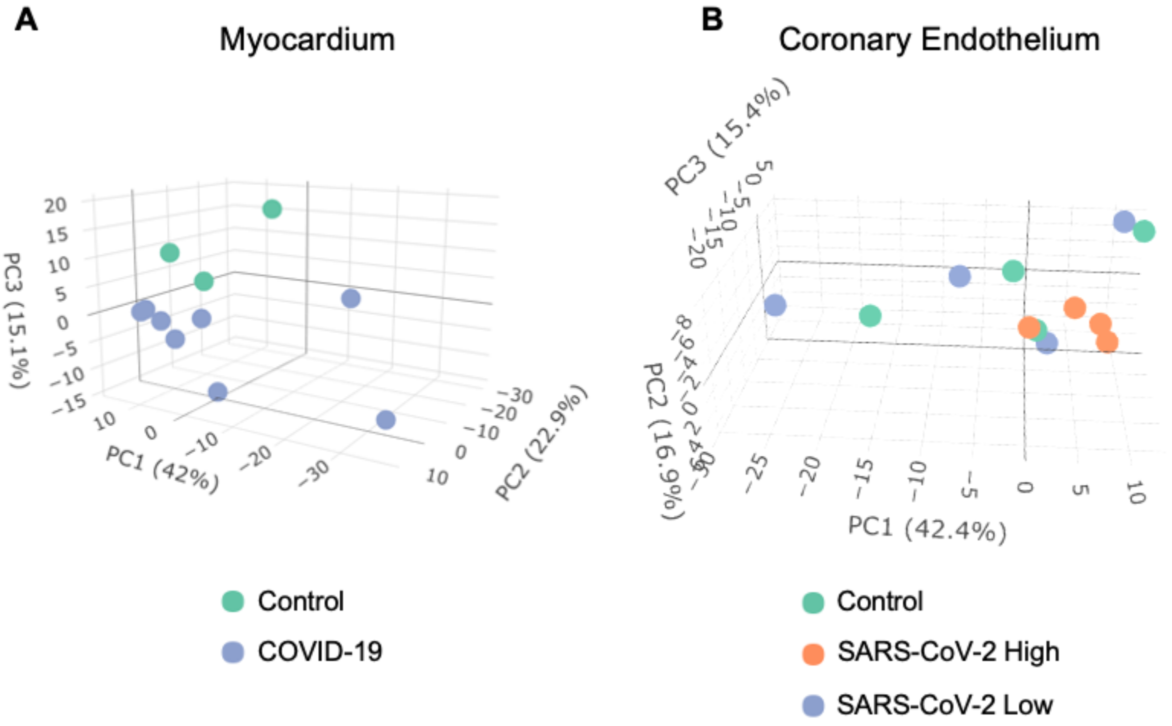


**Figure S4**. Per patient analysis of spatial transcriptomic data does not influence pattern clustering by Principal Component Analysis (PCA). PCA of unsupervised data from myocardial (A) and endothelial (B) samples were analyzed per patient (as opposed to per ROI) as previously described. Importantly, our original findings were unchanged as myocardial samples continued to show pattern clustering, while no association between SARS-CoV-2 and the endothelium was detected. 1454 and 1302 genes were above the limit of quantification, respectively.

**Data Set**

Anonymized datasets generated during this study will be available on **Mendeley Data**.

### Major Resources Table

| REAGENT OR RESOURCE | SOURCE | IDENTIFIER |
|---|---|---|
| ***Antibodies*** | | |
| SARS-CoV-2 nucleocapsid | GeneTex | GTX135361 |
| CD31 | Abcam | Ab9498 |
| ***Biological samples*** | | |
| Patient paraffin embedded myocardial tissue | UAB Tissue Biorepository Core Facility | https://sites.uab.edu/tissuebank/ |
| ***Chemicals, peptides, and recombinant proteins*** | | |
| Xylene | Fisher Scientific | X5P-1GAL |
| 100% Ethanol | Fisher Scientific | HC-800-1GL |
| 95% Ethanol | Fisher Scientific | HC-1100-1GL |
| Tris-EDTA ph9 buffer | Abcam | ab93684 |
| Proteinase K | Fisher Scientific | 25-530-049 |
| Glycine | Sigma-Aldrich | G7126 |
| Elution buffer | Teknova | T1485 |
| Neutral Buffer Formalin 10% | EMS diasum | 15740-04 |
| Ethanol 200 proof | Fisher Scientific | BP2818500 |
| 20X SCC | Sigma-Aldrich | S6639-1L |
| DEPC water | Fisher Scientific | 4387937 |
| Agencourt AMPure XP | Beckman-Coulter | A63880 |
| 10% Tween-20 | Teknova | T0710 |
| Formamide | Fisher Scientific | AM9342 |
| Bovine Serum Albumin Fraction V | Fisher Scientific | BP1605-100 |
| R-Phycoerythrin (R-PE), lightning-link conjugation kit | Abcam | ab102918 |
| Alexa Fluor 488, lightning-link conjugation kit | Abcam | ab236553 |
| DAPI | Biolegend | 422801 |
| Syto61 | Thermofisher | S11343 |
| ProLong Gold Antifade mounting media | Thermofisher | P36934 |
| ***Critical commercial assays*** | | |
| GeoMx Cancer Transcriptome Atlas (COVID-19) | Nanostring | https://www.nanostring.com/products/geomx-digital-spatial-profiler/geomx-rna-assays/geomx-cancer-transcriptome-atlas/ |

| | | |
|---|---|---|
| ***Software and algorithms*** | | |
| Prism v9 | GraphPad | https://www.graphpad.com/scientific-software/prism/ |
| R v3.5.2 | R project | https://www.r-project.org/ |
| Microsoft Excel | Microsoft | https://www.microsoft.com/en-us/microsoft-365/p/excel/cfq7ttc0k7dx?activetab=pivot:overviewtab |
| GeoMx DSP data center | Nanostring | https://www.nanostring.com/products/geomx-digital-spatial-profiler/geomx-data-center/ |
| Basespace | Illumina | https://login.illumina.com/platform-services-manager/?rURL=https://basespace.illumina.com&clientId=basespace&clientVars=aHR0cHM6Ly9iYXNlc3BhY2UuaWxsdW1pbmEuY29tL2Rhc2hib2FyZA&redirectMethod=GET#/ |

### Supplemental References

1. Leek JT, Johnson WE, Parker HS, Jaffe AE, Storey JD. The sva package for removing batch effects and other unwanted variation in high-throughput experiments. *Bioinformatics*. 2012;28:882-883

2. Johnson WE, Li C, Rabinovic A. Adjusting batch effects in microarray expression data using empirical bayes methods. *Biostatistics*. 2007;8:118-127
